# The diagnostic value of RASSF1A promoter methylation in prostate cancer: A systematic review and meta-analysis

**DOI:** 10.1101/2020.07.15.204024

**Authors:** Su-Liang Li, Ye-Xing Li, Yun Ye, Xiao-Hua Yuan, Jian-Jun Wang, Hong-Lin Yan

**Affiliations:** Department of Laboratory Medicine, The First Affiliated Hospital of Xi’an Medical University, Xi’an, Shaanxi 710077, China; An De college, Xi’an University of Architecture and Technology, Xi’an, Shaanxi 710311, China; Emergency Department, The First Affiliated Hospital of Xi’an Medical University, Xi’an, Shaanxi 710077, China; Gastroenterology Department, The First Affiliated Hospital of Xi’an Medical University, Xi’an, Shaanxi 710077, China

**Keywords:** RASSF1A promoter, methylation, prostate cancer, meta-analysis

## Abstract

**Background:** RASSF1A promoter methylation is consistent with clinicopathological data and has good accuracy in distinguishing tumors. However, the diagnostic parameters vary among previous studies. A systematic review was conducted to explore the diagnostic value of RASSF1A promoter methylation in prostate cancer.

**Methods:** A comprehensive search of the literature in the PubMed, Medline, Cochrane Library, Embase and ISI Web of Science databases up to May 21, 2020 was performed. STATA software version 12.0 and Meta-disc version 1.4 were used to analyze the data.

**Results:** The pooled sensitivity was 0.64 (95% CI 0.61–0.66), the pooled specificity was 0.80 (95% CI 0.77–0.83), the PLR was 3.82 (95% CI 1.96–7.44), and the NLR was 0.29 (95% CI 0.16–0.52). Furthermore, the pooled DOR of RASSF1A promoter methylation for prostate cancer was 13.08 (95% CI: 6.56–26.08). The area under the summary ROC curve was 0.87 (95% CI: 0.84–0.90). The results of the meta-regression suggested that heterogeneity was mainly derived from publication year. Fagan’s nomogram showed that the predictive accuracy was increased significantly by detecting RASSF1A promoter methylation for diagnosing prostate cancer.

**Conclusion:** This meta-analysis suggests that detection of the RASSF1A promoter methylation status can be used for the diagnosis of PCa. In the future, further analyses and studies of larger sample sizes in large centers are needed to confirm our conclusion.

## 1. Introduction

Prostate cancer (PCa) is one of the most common malignant tumors of the male urinary system. An estimated 33,330 men are expected to die of the disease in the USA in 2020, and mortality due to PCa accounts for 10% of all cancer deaths.^[1]^ The clinical use of serum PSA and DRE for the early screening and disease progression monitoring of PCa has improved the detection rate of PCa.^[2,3]^ However, PSA and DRE have difficulty meeting clinical needs due to their poor sensitivity or specificity.^[4,5]^ At present, the main problem in the clinical management of prostate cancer is the lack of reliable biomarkers for the early detection and prognosis monitoring of the disease.

The development and impact of epigenetics have been rapidly increasing in recent years.^[6]^ DNA methylation is one of the key phenomena of epigenetics and one of the earliest gene modification methods discovered so far, generally occurring at the CpG site.^[7]^ It plays an important role in the occurrence and growth of many human solid tumors, including prostate cancer. ^[8]^ Hypermethylation of tumor suppressor genes (TSGs) and inactivation of TSG expression are important markers of tumor generation and growth, and the CpG island located in the promoter region of the RASSF1A gene is the most prone to methylation. ^[9]^ A previous study found that RASSF1A promoter methylation was consistent with the clinicopathological data and had good accuracy in distinguishing tumors.^[10]^ However, the diagnostic parameters vary among previous studies. Therefore, a systematic review was conducted to explore the diagnostic value of RASSF1A promoter methylation in prostate cancer.

## 2. Methods

### 2.1. Search strategy

A meta-analysis was performed in accordance with the guidelines of the Preferred Reporting Items for Systematic Reviews and Meta-Analyses (PRISMA).^[11]^ A comprehensive search of the literature in the PubMed, Medline, Cochrane Library, Embase and ISI Web of Science databases up to May 21, 2020 was performed. The following search strategy was used: (“Ras association domain family 1” OR “RASSF1A”) and (“Methylation” OR “Methylations” OR “Hypermethylation”) and (“prostate cancer” OR “prostate carcinoma” OR “prostate neoplasm” OR “prostate tumor”). Subsequently, eligible studies were included for further screening.

### Study selection

The literature search and study selection were independently performed by two researchers (Li and Wang). Any disagreements were resolved by group discussion until a consensus was reached. Studies were included if they met the following study inclusion criteria: (1) Assessment of the methylation of RASSF1A in the study of patients who were diagnosed with prostate cancer; (2) a definite pathological diagnosis of prostate cancer; (3) case-control studies containing at least two comparison groups, with the control group involved patients with benign prostate disease or healthy people; and (4) explicit mention of sensitivity, specificity and critical values in the study. The exclusion criteria were as follows: (1) studies that lacked a control group, only including the case group with a pathological diagnosis of PCa; (2) reviews, comments, meta-analysis, case reports and articles with an indefinite diagnostic threshold; (3) cell lines or animal experiments; and (4) duplicate records.

### Data extraction

The data were collected from the included studies, and the following items were extracted: first author, publication year, country, number of samples, ages, detection method for RASSF1A methylation, sample type, and TP, FP, FN, TN of RASSF1A methylation in diagnosing prostate cancer.

### Quality assessment

The quality of the included studies was evaluated in accordance with the Quality Assessment of Diagnostic Accuracy Studies (QUADAS-2) tool in this meta-analysis. The two authors (Ye and Li) independently completed the quality evaluation.^[12]^ All authors had previously agreed to consider the final determinant of the literature. This study did not require ethical approval or patient written informed consent because it was a systematic review and meta-analysis.

### Statistical analysis

RevMan 5.3 was used to perform the quality assessment, and Meta-disc version 1.4 (version 1.4; Ramón y Cajal Hospital, Madrid, Spain) and the statistical software Stata version 12.0 (Stata Corp LP, College Station, TX, USA) were used to conduct other analyses. The sensitivity, specificity, PLR, NLR, DOR and corresponding 95% confidence intervals (CIs) were calculated from the TP, FP, FN, and TN values, which were extracted from each study before data pooling. We applied a bivariate random effects model^[13]^ to summarize the sensitivity, specificity, PLR, and NLR, and used a hierarchical regression model to summarize the summary receiver operating characteristic (sROC) curve and the area under the ROC curve.^[14]^ The Q statistic and I^2^ were used to assess the statistical heterogeneity across the eligible studies (*P*-values ≤ 0.05 and *I*^2^-values ≥ 50% indicated heterogeneity for the Q statistic). ^[15]^ We conducted meta-regression analyses on the basis of age, publication year, race, detection method and sample type.^[16]^ Deeks’ asymmetry test was used to evaluate potential publication biases, ^[17]^ and Fagan’s nomogram was used to evaluate the pretest probability and posttest probability of the PLR and NLR.^[18]^ A *P*-value < 0.05 was considered statistically significant, and all tests were two-tailed.

## Results

### Literature search

According to the retrieval strategy and inclusion criteria, a total of 206 references were selected from the database; after 96 repeated studies were excluded, 110 references remained. After reading the literature titles and abstracts, 59 studies with outcome observation indicators inconsistent with the purpose of this meta-analysis were excluded, and a total of 15 studies consisting of letters, reviews, comments, meta-analyses and case reports were excluded. Additionally, 16 studies were excluded because they were animal or cell line studies, had index details missing or were not case-control studies. Finally, a total of 20 studies were included in the present meta-analysis. The results of the selection process are shown in Fig 1.

**Fig 1.**
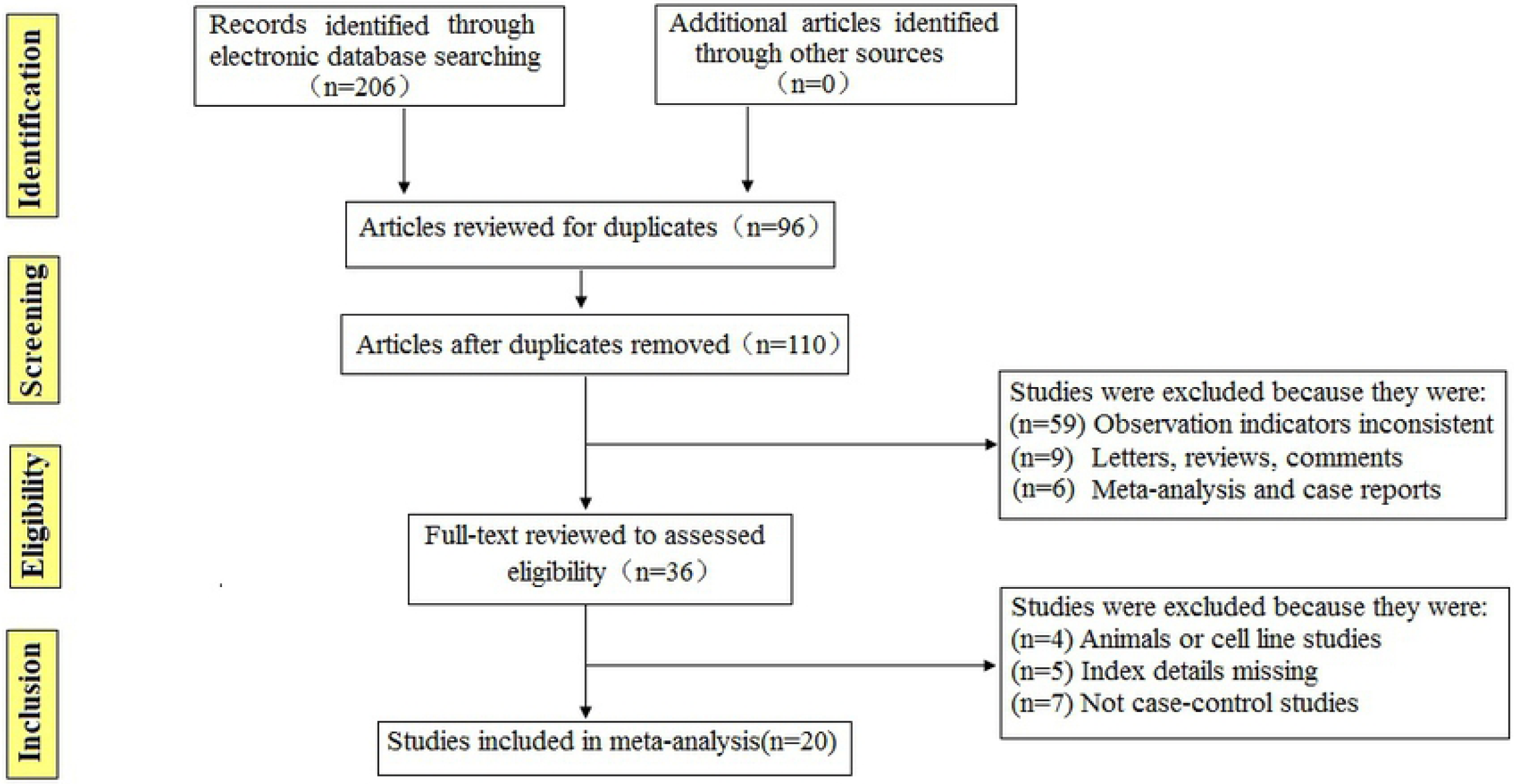
Flow chart showing the study selection procedure.

### Study characteristics

The features of the enrolled studies are listed in Table 1. Twenty studies were published between 2002 and 2018 and included 1640 prostate cancer patients and 769 controls. Eight studies were conducted in Europe, ^[19–26]^ six studies were conducted in Asia,^[27-32]^ and six studies were conducted in North America. ^[33-38]^ The quality assessment results of the included studies showed that the included studies had high quality and could be used for meta-analysis. The results are shown in Fig 2.

**Fig 2.**
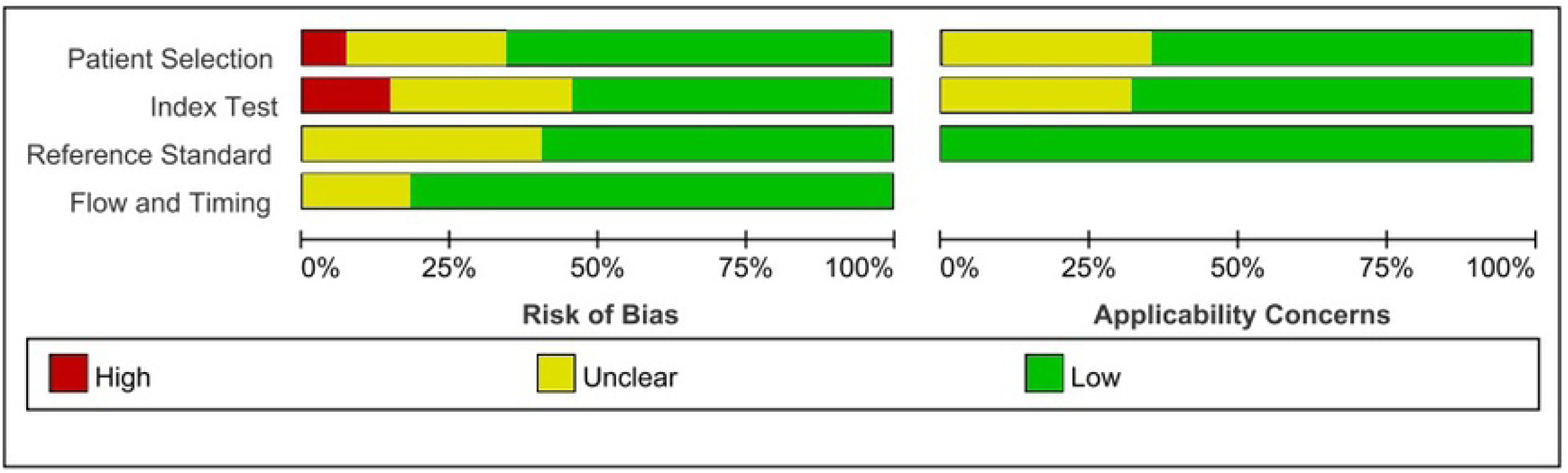
Quality assessment of the included studies.

### Meta-analysis

The summary results for sensitivity, specificity, PLR and NLR are presented in Fig 3. The pooled sensitivity was 0.64 (95% CI 0.61–0.66), the specificity was 0.80 (95% CI 0.77–0.83), the PLR was 3.82 (95% CI 1.96–7.44), and the NLR was 0.29 (95% CI 0.16–0.52). Furthermore, we noted that the pooled DOR of RASSF1A promoter methylation for prostate cancer was 13.08 (95% CI: 6.56–26.08) (Fig 4). Finally, the area under the summary ROC curve was 0.87 (95% CI: 0.84–0.90) (Fig 5).

**Fig 3.**
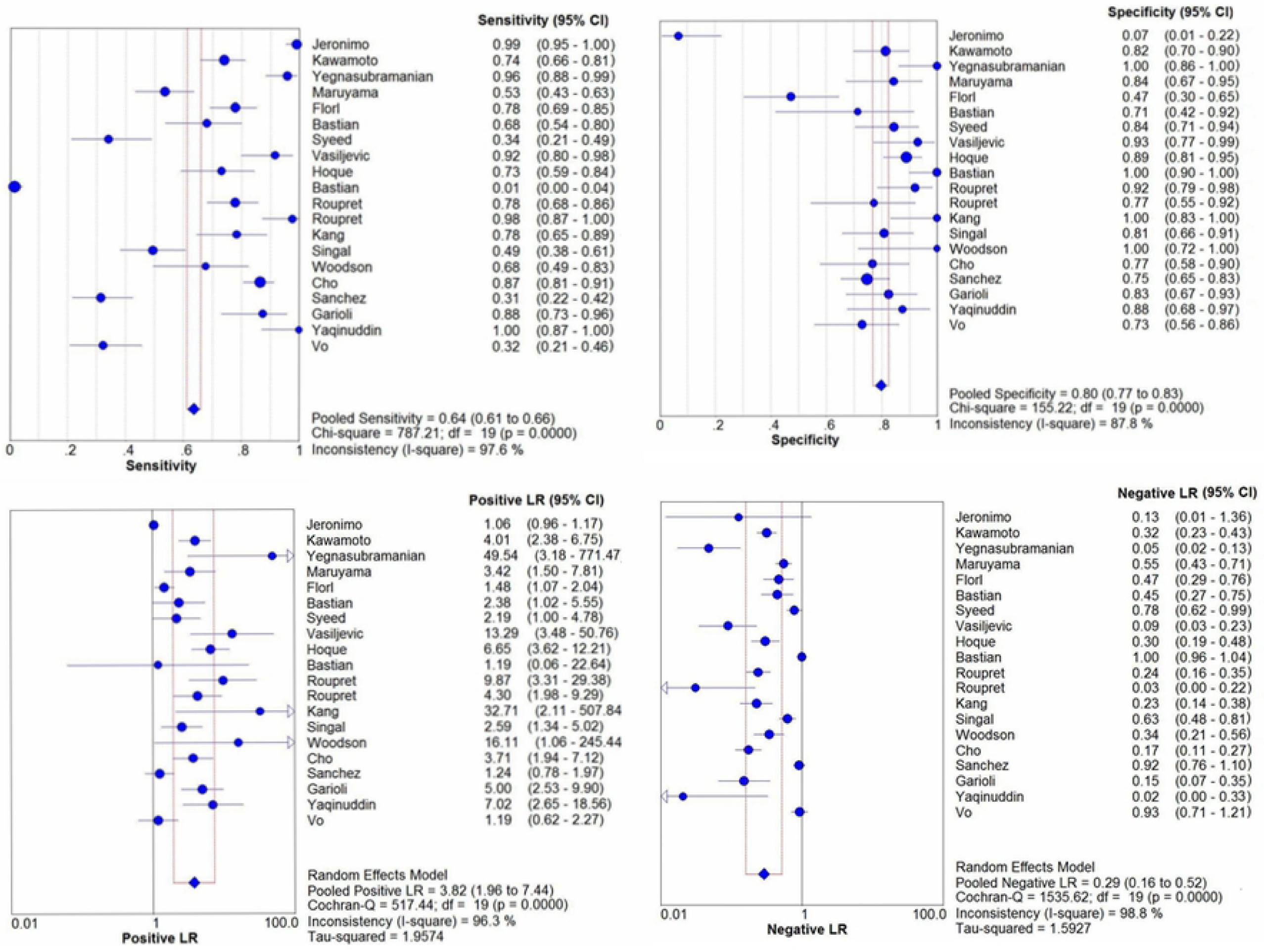
Forest plots for sensitivity, specificity, PLR and NLR. PLR=positive likelihood ratio, NLR= negative likelihood ratio.

**Fig 4.**
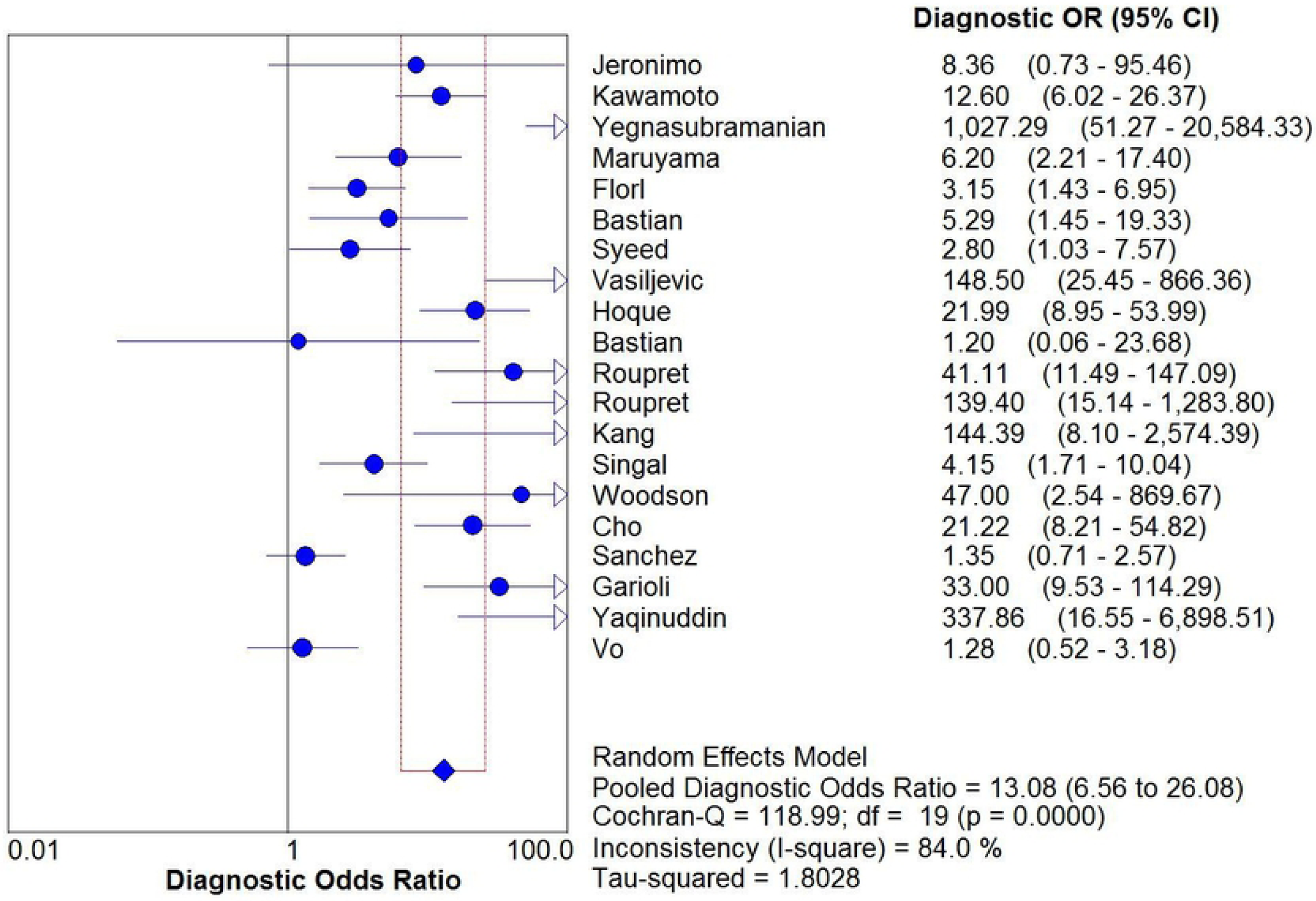
Forest plot for DOR. DOR = Diagnostic odds ratio.

**Fig 5.**
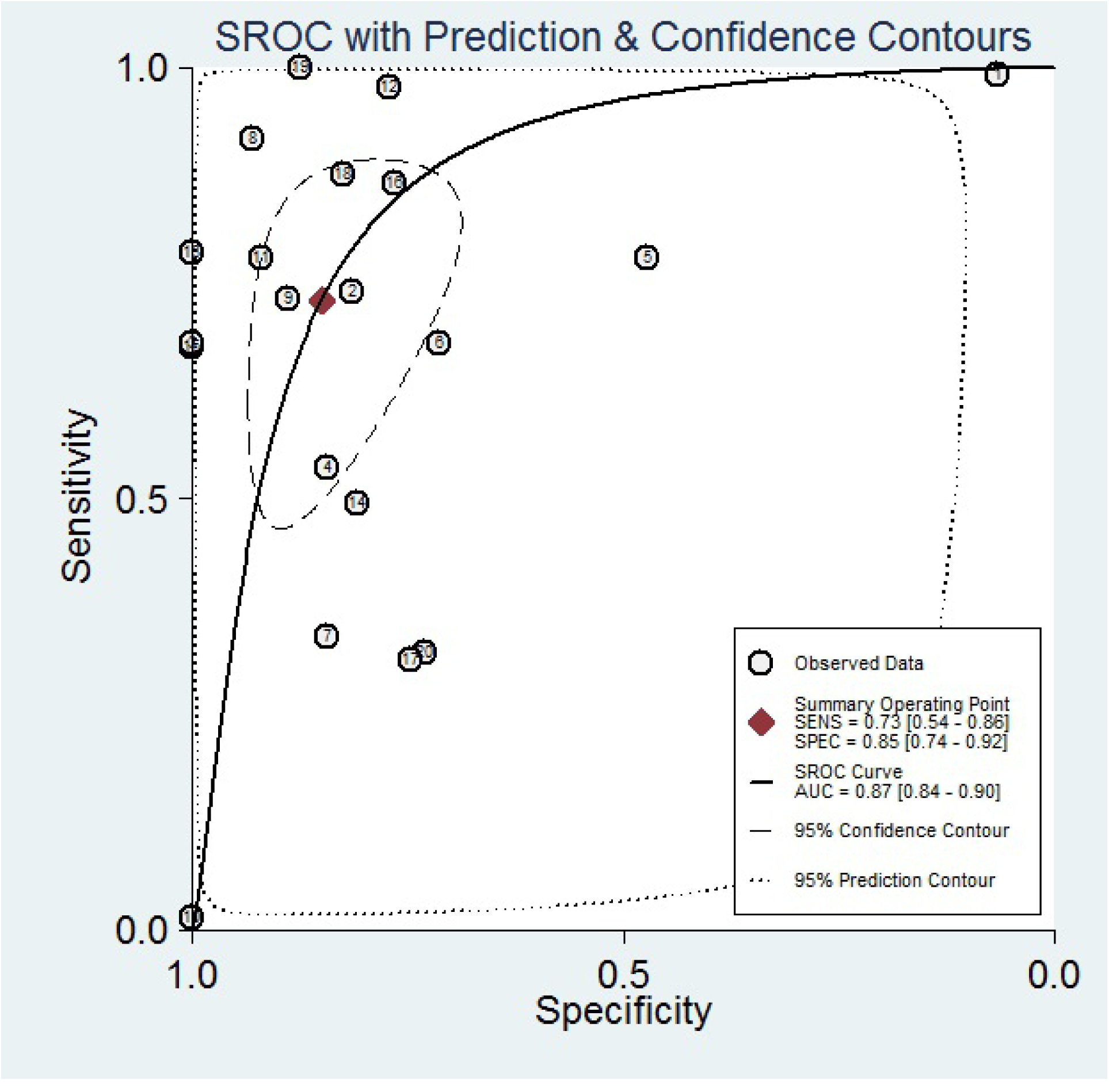
sROC curve of RASSF1A promoter methylation for the diagnosis of prostate cancer in all studies. sROC= summary receiver operating characteristic.

### Heterogeneity analysis

The pooled DOR was 13.08, with significant heterogeneity (*I*^2^=84.0%, *P*≤0.05), and meta-regression was conducted based on age, publication year, race, detection method and sample type. The results suggested that heterogeneity was mainly derived from publication year (Fig 6) (Table 2).

**Fig 6.**
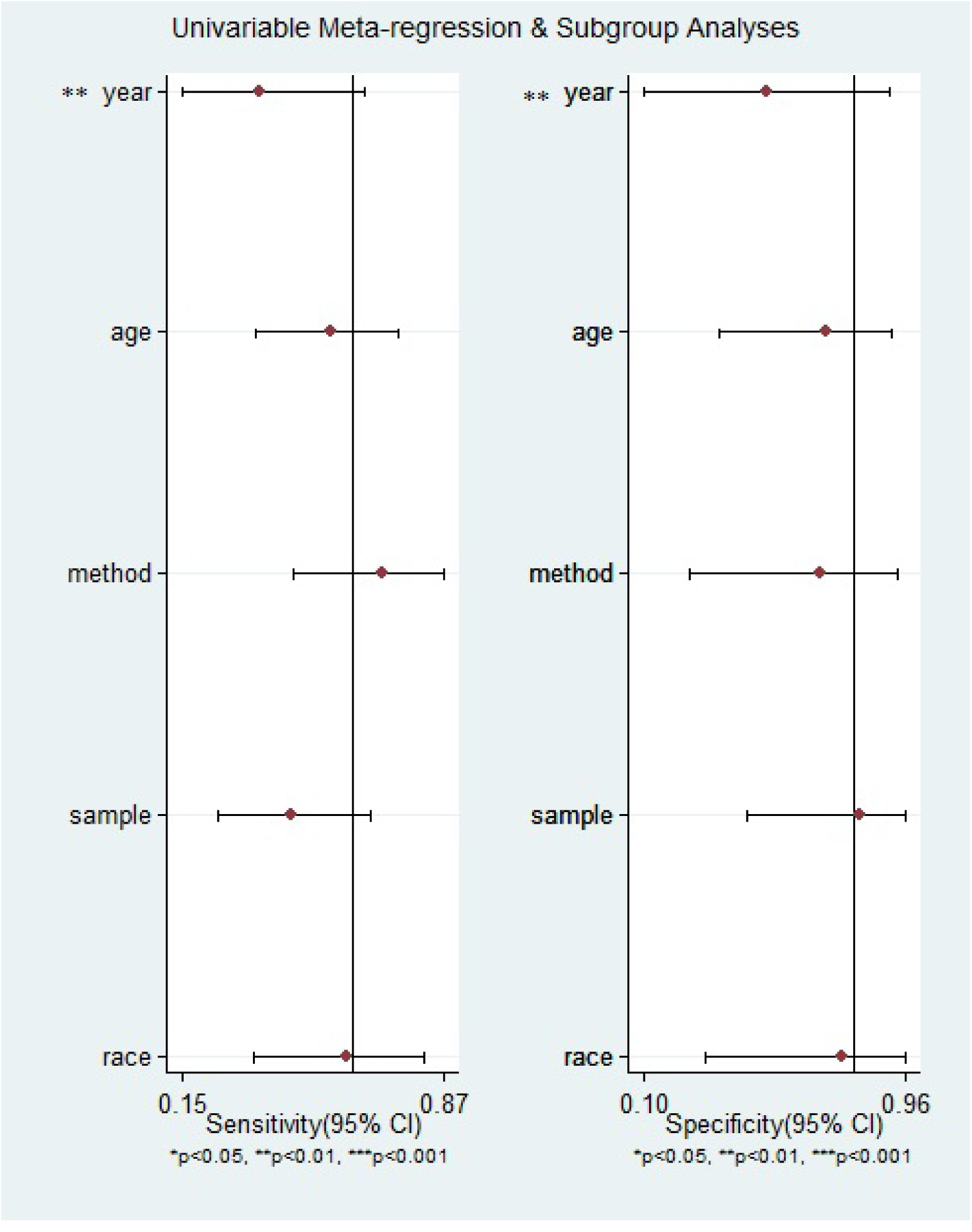
Meta-regression analyses for DOR. DOR = Diagnostic odds ratio.

### Clinical diagnostic efficiency

The changes in the pretest probability and the posttest probability in the diagnosis of prostate cancer with RASSF1A promoter methylation were evaluated by analysis of Fagan’s nomogram. The pretest probability of the PLR was 20%, and the posttest probability was 55%. The pretest probability of the NLR was 20%, and the posttest probability decreased to 7% (Fig 7).

**Fig 7.**
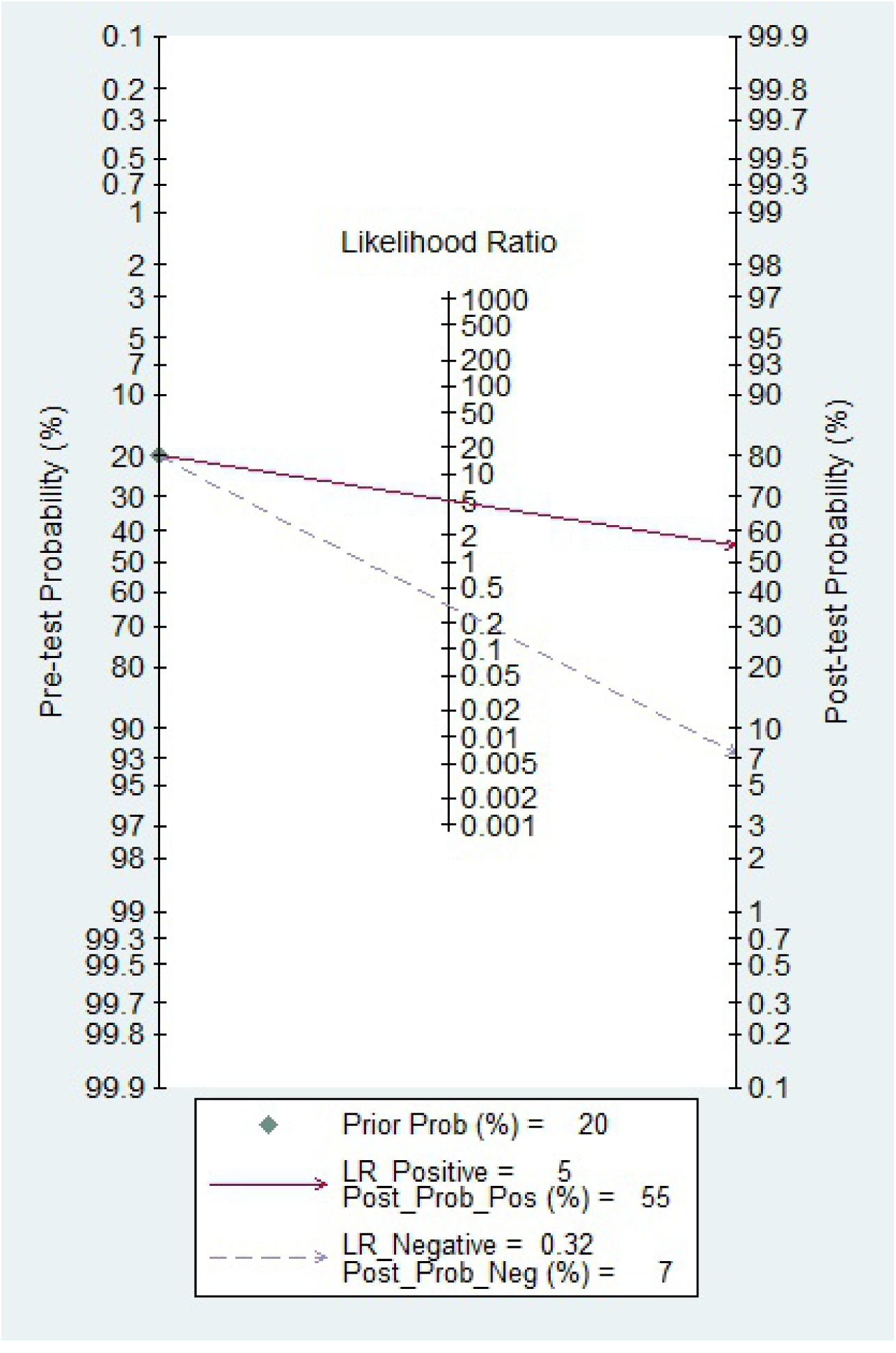
Fagan’s nomogram.

### Publication bias

Deeks’ funnel plot asymmetry test was used to evaluate publication bias. The results of the test showed no evidence of publication bias (*P*=0.73), and the funnel plots were symmetrical (Fig 8).

**Fig 8.**
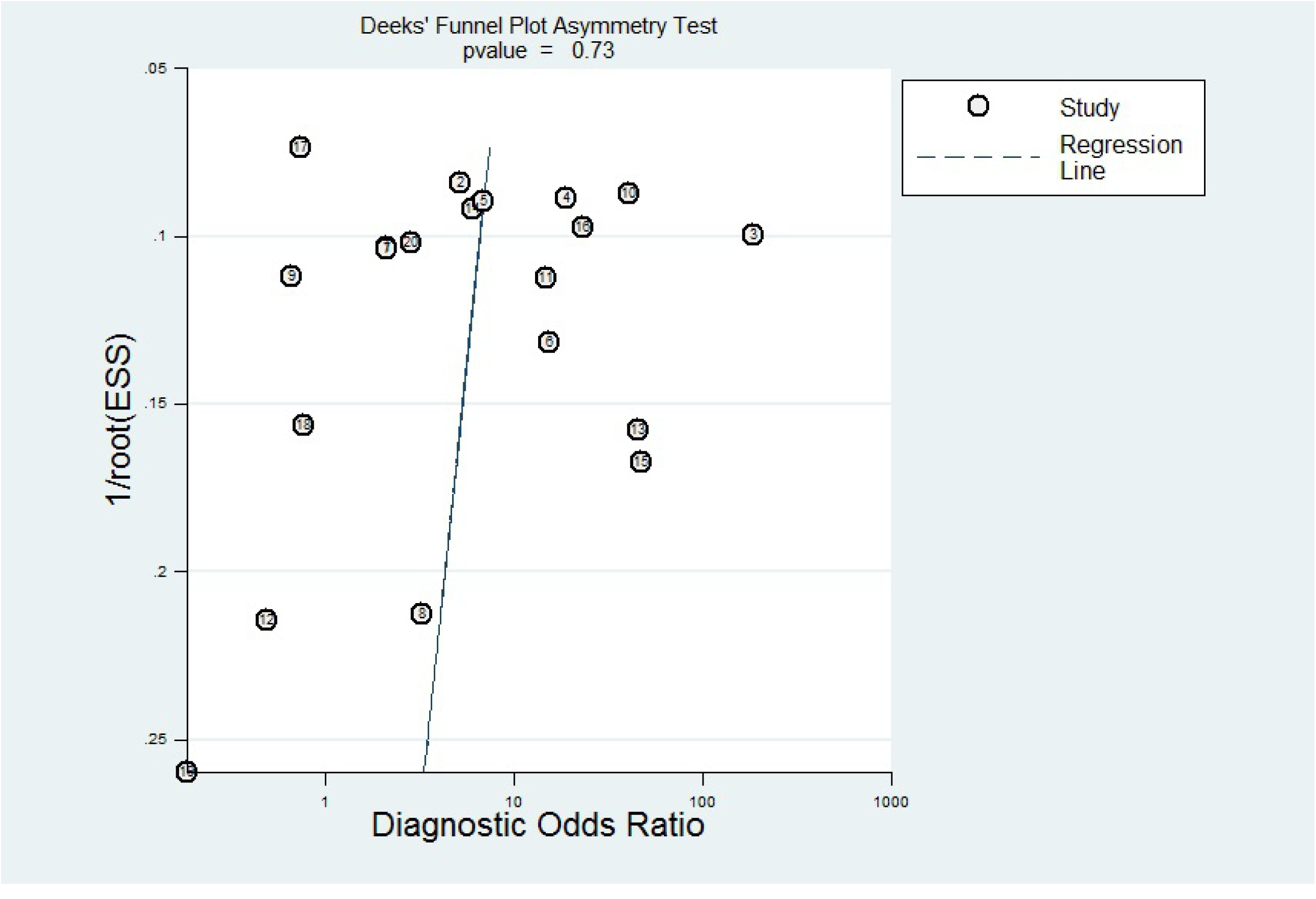
Publication bias.

## Discussion

The American Cancer Society reports that the estimated number of new cases of prostate cancer was 191,930, accounting for 21% of all cases in 2020. ^[1]^ With the increase in life expectancy, the change in dietary structure and the continuous improvement of diagnostic techniques in China, the incidence of PCa has increased rapidly and become the leading threat to men’s health. ^[39]^ PCa is usually asymptomatic in the early stage, but fifty percent of men with prostate cancer die within 30 to 35 months of diagnosis. The early screening of prostate cancer can aid in the early diagnosis and treatment of the disease, ultimately benefitting patients.^[40]^

At present, the main clinical challenge for the diagnosis and treatment of prostate cancer is the lack of reliable biomarkers for the early detection and prognosis monitoring of PCa. ^[41]^ Studies have shown that abnormal DNA methylation plays an important role in the development of tumors, leading to the silencing and activation of some tumor suppressor and protective genes. ^[42]^ Compared with the traditional detection method, RASSF1A promoter methylation has shown better sensitivity and specificity in the diagnosis of prostate cancer. ^[43]^

The meta-analysis investigated the diagnostic value of RASSF1A methylation for prostate cancer and included 20 studies involving a total of 2409 patients. The results show that the pooled sensitivity and specificity were 0.64 and 0.80, with missed diagnosis rates was 0.36 and 0.20, respectively. This result shows that the diagnostic efficiency was high. In this study, the pooled PLR and NLR of RASSF1A methylation for diagnosing prostate cancer were 3.82 and 0.29, respectively, and the results show an acceptable detection rate. These results suggest that the overall accuracy of prostate cancer detection by RASSF1A promoter methylation is relatively good. Next, the pooled diagnostic odds ratio was 13.08, suggesting that RASSF1A methylation has outstanding discrimination ability for prostate cancer. Furthermore, the area under the summary ROC curve was 0.87 (95% CI: 0.84–0.90), indicating high diagnostic value.

However, the results show high heterogeneity with an *I*^2^ value for the heterogeneity test of DOR of 84.0%. Therefore, we used meta-regression analysis to explore the possible sources of heterogeneity. The *P*-value of publication year was 0.01 (<0.05), and the remaining *P*-values were all >0.05. The results suggest that the publication year of the research was the main source of heterogeneity. Deeks’ funnel plot asymmetry test showed no evidence of publication bias (P=0.73).

Fagan’s nomogram showed that the predictive accuracy was increased significantly by detecting RASSF1A promoter methylation for diagnosing prostate cancer. RASSF1A promoter methylation is sensitive to the diagnosis of prostate cancer. The pretest probability of the PLR increased from 20% to a posttest probability of 55%. Furthermore, the absence of RASSF1A promoter methylation reduced the possibility of prostate cancer diagnosis. The pretest probability of the NLR decreased from 20% to a posttest probability of 7%. Dammann found hypermethylation of the RASSF1A promoter in lung cancer for the first time, which was positively correlated with a decrease in gene expression.^[44]^ Studies have shown that RASSF1A promoter methylation is 99.15% sensitive to prostate cancer detection. ^[45]^

This study also has the following limitations: (1) This study only included published literature, so a potential publication bias cannot be excluded. (2) The language of the literature retrieved was limited to English, and relevant studies in other languages may have been missed. (3) Some of the literature included a small number of subjects and did not use blinding methods.

Based on the results of the current meta-analysis, we believe that the detection of RASSF1A methylation can be used as an important method to screen benign and malignant prostate diseases. RASSF1A methylation has broad application prospects in PCa diagnosis, treatment and prevention.

## Conclusion

This meta-analysis suggests that detection of the methylation status detection of the RASSF1A promoter can be used for the diagnosis of PCa. In the future, further analyses and studies of larger samples in large centers are needed to confirm our conclusion.

## Abbreviations

PSA: prostate-specific antigen
DRE: digital rectal examination
TN: true negative
TP: true positive
FN: false negative
FP: false positive
NLR: negative likelihood ratio
PLR: positive likelihood ratio
DOR: diagnostic odds ratio
SROC: summary receiver operating characteristic
AUC: area under the SROC curve
CI: confidence interval
QUADAS-2: Quality Assessment Of Diagnostic Accuracy Studies tool-2
P: P-value of overall effect.

## Author contributions

Data curation: Yun Ye, Xiao-Hua Yuan.

Formal analysis: Su-Liang Li.

Investigation: Ye-Xing Li, Jian-Jun Wang.

Methodology: Ye-Xing Li, Yun Ye.

Software: Su-Liang Li, Yun Ye.

Supervision: Hong-Lin Yan.

Validation: Ye-Xing Li, Su-Liang Li.

Writing – original draft: Yun Ye.

Writing – review and editing: Yun Ye, Su-Liang Li.

## Acknowledgments

This study was supported by the Education Department of the Shaanxi Provincial Government (2020JM-607) and (2017JM-8120). The authors express their gratitude to the study participants and research personnel for their involvement in the study. The authors declare that there are no conflicts of competing interests regarding the publication of this article.

